# The osteoclast intracellular environment fosters bacterial growth during *Staphylococcus aureus* infection

**DOI:** 10.64898/2026.03.18.712565

**Authors:** Luke D O’Connor, Saumya Bhagat, Nidhi Roghati, Gabriel Mbalaviele, James E Cassat, Deborah J Veis

## Abstract

Bone infections, which are predominantly caused by *Staphylococcus (S.) aureus*, can be difficult to treat and have high rates of chronicity and reoccurrence. We previously identified that osteoclasts, the cells that break down bone matrix, may contribute to disease progression by allowing *S. aureus* to replicate intracellularly. There we identified that this bacterium’s ability to grow intracellularly is tied to the maturation of osteoclasts. In this study we addressed whether osteoclast differentiation supports intracellular growth by changing the host cell’s response to infection or by altering the host cell environment to better support *S. aureus*. Using dual species RNA-sequencing we analyzed host and bacterial transcripts of infected osteoclast and precursor bone marrow macrophage (BMM) cultures. Host transcript analysis suggests that infected osteoclasts are slow to upregulate bacterial response genes compared to BMMs. We also identify that the *S. aureus* transcriptional response is primarily determined by the host cell type, and that bacteria in osteoclasts upregulate carbon metabolism genes compared to those inside BMMs. By utilizing intracellular survival assays on *S. aureus* mutants deficient in carbon metabolism and related pathways we determine that *S. aureus* require glycolysis, acetyl-CoA synthesis, and aspartate biosynthesis for proliferation inside osteoclasts, although bacteria can survive without them. With differentiation, osteoclasts increase glutamine uptake, and this metabolite is required for S. aureus intracellular growth. Taken together, these findings suggest that osteoclasts support *S. aureus* intracellular growth by providing nutrients required to replicate in the context of a blunted antimicrobial response.

**IMPORTANCE:** Infectious osteomyelitis, bone infection, is frequently caused by the bacterium *Staphylococcus aureus*. Intracellular infection of cells that build bone, osteoblasts, and cells that resorb bone, osteoclasts, have been implicated in disease progression by providing a niche for immune evasion. While *S. aureus* in osteoblasts are largely quiescent, bacteria in osteoclasts proliferate and therefore may be a source of reemergent infection. Factors that promote this growth in osteoclasts are poorly characterized. In this study we find that osteoclasts have a diminished transcriptional response to infection and show that *S. aureus* acquire glucose and glutamine, which have high flux in osteoclasts, to support intracellular growth. We further observe that *S. aureus* in osteoclasts require aspartate synthesis to grow intracellularly. This work highlights the importance of host cellular metabolism for the intracellular fate of *S. aureus* as an added factor beyond the direct antimicrobial response.

## INTRODUCTION

*Staphylococcus (S.) aureus* driven osteomyelitis is a debilitating disease that is difficult to resolve once established. Staphylococcal Abscess Communities (SACs) and biofilms are considered primary causes of treatment recalcitrance. In both cases extracellular bacteria use physical barriers to immune cell access and increase their tolerance of antibiotics (1–3). *S. aureus* can also infect and survive inside many cells in the bone and bone marrow, but the mechanisms involved are not fully understood (1, 4–7). Osteoclasts (OCs) appear to be a unique reservoir for *S. aureus* in the bone as they allow bacteria to rapidly expand inside of them, providing protection from antibiotics and immune cells while increasing the tissue’s bacterial burden (8, 9). OCs are multinucleated, myeloid derived, cells that resorb mineralized bone matrix (10). Historically OCs were thought too short-lived, a few days to weeks, to be a significant reservoir for intracellular bacteria (11–15). However, *in vivo* lineage tracing showed that OCs live as long as 6 months and can add or subtract nuclei via fusion/fission events (16, 17). These features position OCs as a sufficiently long-lived permissive cell type that allows intracellular bacterial growth to maintain chronic infection.

During OC differentiation, cells undergo numerous changes that impact signaling and metabolism. For instance, when the osteoclastogenic Receptor Activator of NF-κB (RANK) is stimulated by its ligand, RANKL, it induces expression of NFATc1, which alters downstream NF-κB signaling and transcriptional targets by partner switching with AP-1(10). This change turns otherwise inflammatory signals (e.g. TNFα, IL1β, or TLR2 activation) to induce OC gene expression. As a result, inflammation results in increased OC differentiation (7, 18). As mononuclear OC lineage precursors mature, they form large polykaryotic cells by fusing, while increasing their metabolic activity via aerobic and anaerobic respiration (19, 20). To support maturation, they acquire large amounts of iron, glucose, and glutamine (Gln) which are for energy, to build biomass, and to support mitochondrial biogenesis (21–27). Gln is thought to be particularly important due to its metabolism into α-ketoglutarate (α-KG) and other metabolites via glutamate (Glu) (22, 26, 28). These iron, glucose, and Gln derived nutrients are needed to support the energy-intensive function of resorbing mineralized matrix (21–27).

In addition to allowing intracellular *S. aureus* growth, we found that OC maturity plays a role in the magnitude of the bacterial burden increase, with significant increases between mononuclear preosteoclasts (preOCs) to mature multinucleated OCs (mOCs) *in vitro* (8). Furthermore, inhibiting OC differentiation by knocking down NF-κB pathways prevents bacterial growth. This indicates that changes derived from OC differentiation and maturation create an intracellular environment supportive of *S. aureus* growth. Additionally, the greater growth in mOCs compared to preOCs indicates that factors resulting in bacterial survival and expansion may occur at discrete phases or through discrete processes.

This study focuses primarily on cellular changes during OC differentiation resulting in a high intracellular bacterial burden. We hypothesized that OC differentiation promotes intracellular infection by blunting the OC’s defense response and/or by providing increased bacterial access to key nutrients for growth. We first approached this hypothesis by conducting host and *S. aureus* transcriptomic analysis on a time course of intracellular infections using either OCs or their precursors, bone marrow macrophages (BMMs), as hosts. These data confirmed distinct responses of OCs and BMMs to infection, although most differential gene expression was related to host cell type. Using the bacterial transcriptome, we addressed how *S. aureus* respond to the OC host cell environment. In follow up experiments, we utilized a combination of bacterial mutants and manipulations of nutrients in culture media to understand the resources that bacteria acquire from OCs.

## METHODS

### Materials

Materials used in this study can be found in Table 1.

**Table 1:** Materials.

| Table1: Materials |  |  |
| --- | --- | --- |
| Reagent | Manufacturer | Catalog # |
| **Cell Culture** |  |  |
| αMEM | Sigma-aldrich | M0894 |
| Sodium Bicarbonate | Sigama Aldrich | S5761 |
| αMEM without glutamine | Corning | 15-012-CV |
| L-Glutamine (200nM) | Gibco | 25030081 |
| MEM non-essential amino acids | Gibco | 11140050 |
| Fetal Bovine Serum | Gibco | A5670701 |
| RANKL-GST | Lab stock |  |
| CMG 14-12 supernatant (M-CSF) | Lab stock |  |
| **Bacterial Culture** |  |  |
| Trypticase soy broth | BD | 211768 |
| Trypticase soy broth (dextrose free) | BD | 286220 |
| Trypticase soy agar | BD | 211043 |
| Sodium pyruvate (general cell culture) | Gibco | 11360070 |
| **Antibiotics** |  |  |
| Pen Strep | Gibco | 15140122 |
| Erythromycin | Fischer Scientific | AC227330050 |
| Gentamicin | Gibco | 15750060 |
| **Seahorse assay** |  |  |
| XF DMEM | Agilent | 103575-100 |
| Sodium pyruvate | Agilent | 10-357-8100 |
| Glucose | Agilent | 103577-100 |
| Seahorse XF Mito Stress Test Kit | Agilent | 103015-100 |
| Recombinant M-CSF | Lab stock |  |
| **RNA Extraction and processing** |  |  |
| Trizol reagent | Invitrogen | 15596018 |
| Direct-zol RNA miniprep | Zymo Research | R2072 |
| Rino lysis bead kit(yellow) | Next advance | YELLOWR1-RNA |
| FastSelect Epidemiology Kit | Qiagen | 333272 |
| **Equipment** |  |  |
| Bullet Blender Storm Pro | Next Advance |  |
| Seahorse XFe96 Analyzer | Agilent |  |
| Nanodrop 2000 | Fischer Scientific |  |
| NovaSeq X Plus | Illumina |  |

### Primary cell culture

Primary murine bone marrow derived macrophages were isolated from 9-13 week old C57Bl/6J mice and expanded in macrophage expansion media- αMEM supplemented with 10% heat inactivated (HI) Fetal Bovine Serum, 1% Penicillin/streptomycin, and 1:10 dilution of CMG14-12 supernatant as a source of M-CSF(29). Osteoclasts were differentiated and cultured in αMEM supplemented in with 10% HI-FBS, 1:50 dilution of CMG14-12 supernatant, and 90ng/mL RANKL-GST(30).

### Bacterial strains and culture conditions

Strains used in this study are listed in Table 2. Bacteria were cultured in Trypticase Soy Broth (TSB) unless otherwise specified. The *pyk* mutant was cultured in dextrose-free TSB supplemented with 1% sodium pyruvate. To obtain growth curves, overnight cultures were diluted 1:100 into TSB, modified as above for the *pyk* mutant, and OD600 measured every 30min.

**Table 2:** Bacterial strains.

| Table 2: Bacterial strains |  |  |
| --- | --- | --- |
| Strain | Description | Reference |
| TI3 | USA300 lineage Osteomyelitis clinical isolate | (9) |
| WT LAC | USA300 LAC (AH1263) with Erm sensitivity. Background for most mutants | (76) |
| $\Delta pyk::erm$ LAC | Glycolysis mutant | (41, 69) |
| $pdhA::tn$ LAC | Metabolism mutant | (41, 77) |
| $acnA::tn$ LAC | Metabolism mutant | (41, 77) |
| $sucA::tn$ LAC | Metabolism mutant | (41, 77) |
| $gudB::tn$ LAC | Metabolism mutant | (41, 77) |
| $aspA::tn$ LAC | Metabolism mutant | (41, 77) |
| $alsT::tn$ JE2 | Transport mutant | (77) |
| $gltS::tn$ LAC | Transport mutant | (41, 77) |
| $gltT::tn$ LAC | Transport mutant | (41, 77) |
| $aspA::tn/gltT::tn$ LAC | Metabolism and transport mutant | (41) |
| $aspA::tn$ $pOS1-gltT$ LAC | Metabolism mutant with transport overexpression | (41) |

### Gentamycin protection assay

Overnight bacterial cultures were diluted 1:100 and subcultured to an OD600=1 in TSB, approximating 10^9 CFU/mL. Bacteria were pelleted and resuspended in PBS. Host cells, cultured in antibiotic free media >24hrs, were inoculated with the multiplicity of infection (MOI) and duration indicated below. Extracellular *S. aureus* were eliminated by treatment with 300ng/mL gentamycin for 1hr. Samples for were washed with PBS and either collected or had their antibiotic free media replenished at 1.5 hours post infection (hpi). For experiments with media manipulation, either addition of 2-deoxyglucose or Gln restriction, conditions were introduced during media replenishment at 1.5hpi. Mock infection samples shared the same media changes and gentamycin treatment.

### Dual RNA-sequencing

OCs and BMMs were seeded onto a 6-well plate and inoculated with the clinical isolate, TI3, at a MOI 100 for 1hr prior to the gentamicin protection assay. RNA was extracted using a protocol adapted from (31), which results in host and bacterial RNA enriched fractions. Briefly, murine cells were lysed with TriZol on ice for 5min. Cell lysates were centrifuged at 21,000g for 20min at 4°C, and host RNA-enriched supernatant was separated. The resulting pellet was transferred to a yellow Rino Lysis tube (Next Advance), resuspended in Trizol. Bacteria were mechanically lysed by running a bullet blender for 10 min at max speed. RNA from the bacterial and host fractions was isolated and quantified separately using a Zymo Direct-zol Miniprep kit (Zymo Research) and nanodrop (Thermo Scientific). Samples were reconstituted at a ratio of 85% host-enriched RNA to 15% bacteria-enriched RNA to allow for sufficient read depth of both species. Biological replicates were pooled so that each sequenced sample contained cells from 3 different mice each infected with different inocula. Pooled samples were then submitted to the Genome Technology Access Center at Washington University’s McDonnell Genome Institute for rRNA depletion using the Qiagen FastSelect Epidemiology kit, library preparation, and sequencing to a target depth of 50 million reads on an Illumina NovaSeq X Plus sequencer.

### RNA analysis

Reads were aligned and annotated using the HISAT2-Rsubread-featureCounts-DeSeq2 pipeline(32–36). We used Mm39 (GCA_000001635.9) as our murine reference genome (37). *S. aureus* USA300_FPR3757 (ASM1346v1, GCA_000013465.1) was used as the reference bacterial genome, as done with our previous work (9, 38). Raw datasets required for this analysis are available on the Genome Expression Omnibus at GSE315840. Code for the analyses are available on github.

### Colony Forming Unit (CFU) Assays

Expanded murine BMMs were differentiated until mature OCs were observed (3-5 days) on 12-well plates and inoculated with a MOI10 for 30min prior to the gentamicin protection assay. Intracellular *S. aureus* were harvested by lysing host cells in ice cold water for 10min. Samples at 1.5hpi and 18hpi were quantified by plating sample dilutions on TSA plates. Each condition was run in technical triplicate then averaged. 18hpi well media was sampled for planktonic bacteria; wells with substantial detectible extracellular bacterial growth were omitted from analyses. The average ΔLog_10_(CFU) (abbreviated ΔCFU subsequently) was calculated using the following equation: ΔLog_10_(CFU) = Average[Log_10_(CFU_18hpi_) - Log_10_(CFU_1.5hpi_)]. Experiments were also omitted when OCs were not sufficiently differentiated, as determined by a WT LAC technical control with ΔCFU <0.50. Data points reported in figure panels are biological replicates representing unique combinations of OC cell prep and treatment (mutant strain and/or media manipulation). These biological replicates which share an OC cell prep, and therefore part of the same individual experiment, have been color coded.

For experiments with OC media manipulation, either by addition of 2-deoxyglucose or removal of glutamine, these conditions were applied after 1.5hpi to control for possible effects to OC viability and differentiation or *S.aureus* internalization. As cells were treated identically until 1.5hpi collection, the 1.5hpi CFU quantification was shared among 18hpi CFU of all media conditions within an experiment to generate the ΔCFU.

### Oxygen Consumption Rate Analysis

Methods were adapted from (39). Cells were differentiated into OCs for 3-4days or seeded in macrophage expansion media. Cells were inoculated with TI3 at a MOI 100 for 1hr prior to the gentamicin protection assay. 1hr prior to mitochondrial stress test, media was changed to XF DMEM supplemented with 10mM glucose, 2mM-L-Gln, 10µM sodium pyruvate, and 100ng/mL M-CSF, +/- 90ng/mL RANKL, and equilibrated in a non-CO_2_ incubator. We next ran a Mito stress test according to Agilent specifications, with a 1.5min mixing period and 5min measurement period. OCR and ECAR calculation and analysis was conducted on the Seahorse XFe96 software, Wave 2.6.1.

### Statistical analysis

All intracellular growth analyses were conducted using ΔCFU. For ΔCFU analysis comparing mutant strains to WT LAC, all experimental conditions without media manipulation were combined and analyzed by one-way ANOVA followed by Dunnett’s test, with a family wide α = 0.05. This allowed us to make head-to-head comparison of a given mutant strain and WT LAC while limiting our type I error. Figure panels use p-values generated from this analysis but display data subsets which use samples from the same set of experiments. Analysis of mutant internalization was performed on 1.5hpi CFU quantification data, using a one-way ANOVA. Data from control conditions from media manipulation experiments (i.e. without any manipulation) were included in this internalization analysis (Figure S6B) and the ΔCFU analysis (Figure S6A). Likewise, controls in the Gln restriction experiments (Figure 6D) were also included in these global analyses. Experiments testing the effect of 2-Deoxyglucose were analyzed by one-way ANOVA followed by Dunnett’s test, with family wide α = 0.05. Experiments testing effect of Gln restriction and mutant strains were analyzed by two-way ANOVA.

All statistical analyses noted above were conducted using R (version 4.2.1). Code and documentation of the packages used for these analyses are available on github.

Mitochondrial stress test data were analyzed by two-way ANOVA in GraphPad Prism 10. Conditions were only analyzed within individual experiments as basal oxygen consumption rate varies between cell preparations.

### Reanalysis of metabolomics

Amino acid abundances were reanalyzed using data generated by Hu et al, available in the Texas Data Repository https://doi.org/10.18738/T8/X6CDEQ (22).

## RESULTS

### Host cell environment establishes *S. aureus* transcriptional response

Our previous studies showed that OC differentiation and maturation are key to supporting bacterial growth, yet it is unclear whether OCs are permissive hosts due to changes in their antimicrobial response or the intracellular nutrient availability(8, 9). We first addressed this question by conducting time-course dual RNA-sequencing on BMMs and OCs infected with a clinical *S. aureus* strain, TI3, derived from an osteomyelitis patient and capable of causing osteomyelitis in a murine model (Figure 1A) (9, 40). We have observed that it takes 8-10 hours post infection (hpi) for bacterial load to increase in OCs, but we hypothesized that transcriptional changes occur before then. Therefore, samples were collected prior to infection (0hpi), after intracellular selection (2hpi), at a timepoints prior to and after bacterial growth in OCs (6hpi and 21hpi respectively) (8). Principle component analysis (PCA) of intracellular *S. aureus* transcripts indicates that bacterial samples cluster primarily by their host cell type, with smaller separation based on the collection timepoint (Figure 1B). Data from inocula (0hpi) were omitted to improve sensitivity of downstream analyses (Figure S1).

**Figure 1:**
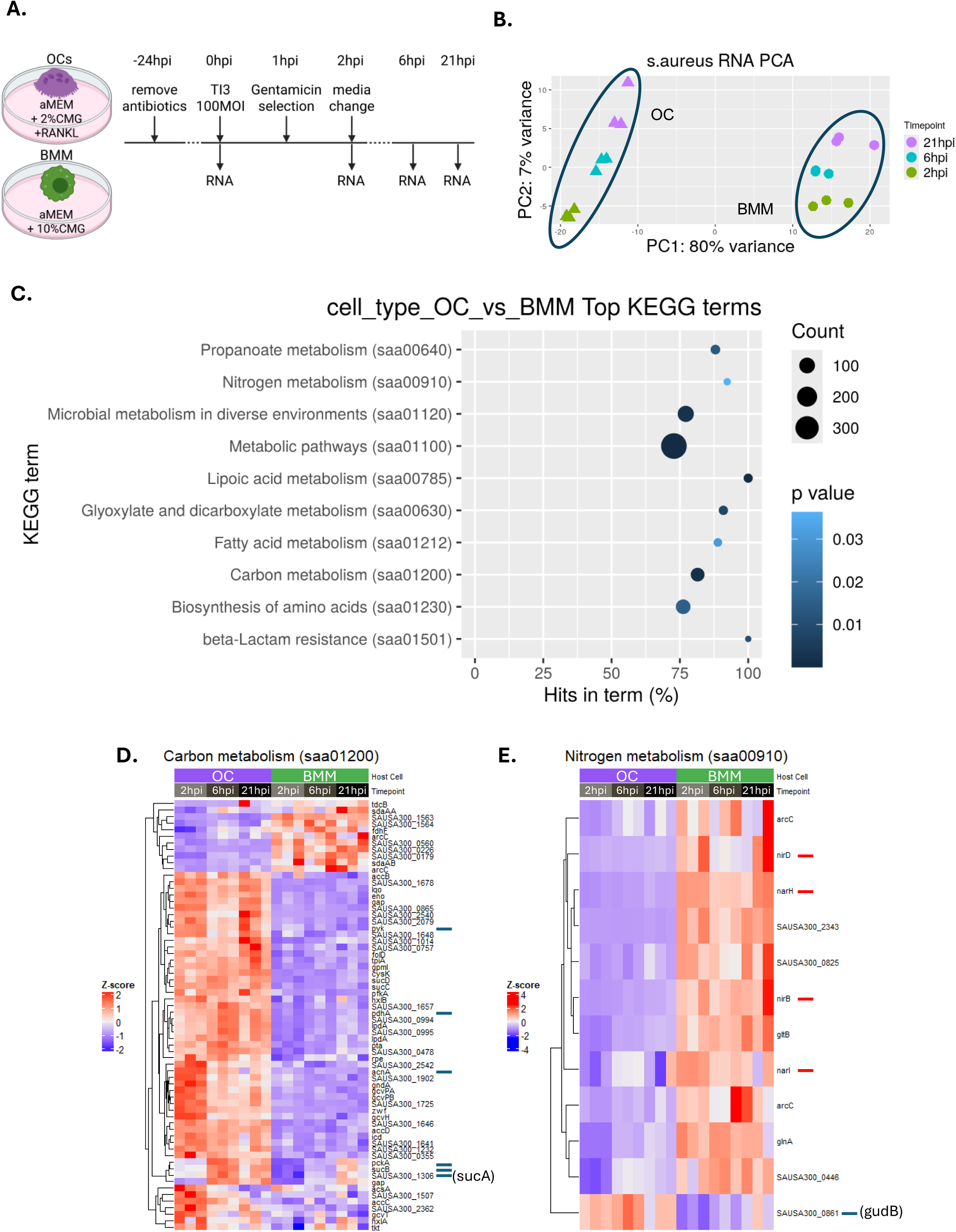
*S. aureus* response to internalization is established by host cell type. **A)** Dual species RNA-seq experimental design. **B)** Principal component analysis of intracellular *S. aureus* RNAseq samples 2-21hpi. Bacterial samples cluster primarily by host cell type. **C)** KEGG pathways enriched for genes differentially expressed by *S. aureus* in OCs compared to BMMs. Heatmaps of *S. aureus* genes in **D)** Carbon metabolism and **E)** Nitrogen metabolism pathways. Z-score represents transcript abundance relative to all transcripts in terms of standard deviation(s). Blue ticks = genes functionally tested by mutant strain, red ticks = genes discussed.

We further investigated the impact of OC versus BMM cellular environment on *S. aureus* transcript profiles by using a Wald analysis to analyze differentially expressed genes (DEGs) based on host cell type. Subsequent KEGG pathway analysis revealed that *S. aureus* differentially express Metabolic Pathways (saa01100), including several subterms (Figure 1C; Figure 2A). Importantly, pathway analysis did not detail which *S. aureus* condition was enriched for these DEGs. Therefore, we examined DEGs within the pathways (heatmaps, Figure 1D-E; Figure S2) and within the context of the KEGG metabolic pathway map (Figure 2). We observe that *S. aureus* in OCs express higher levels of genes related to glycolysis, acetyl-CoA synthesis, gluconeogenesis, fatty acid elongation, the tricarboxylic acid (TCA) cycle, and several amino acid metabolism pathways that feed into the TCA cycle (Figure 2B, purple; Table S1 and Figure S2). Of note, *aspA*, an aspartate transaminase that we previously showed is important during bone infection *in vivo,* was also differentially expressed in our dataset but is not annotated by KEGG (41, 42). Therefore, *aspA* was included in our analysis, despite its absence in our pathway analysis.

**Figure 2:**
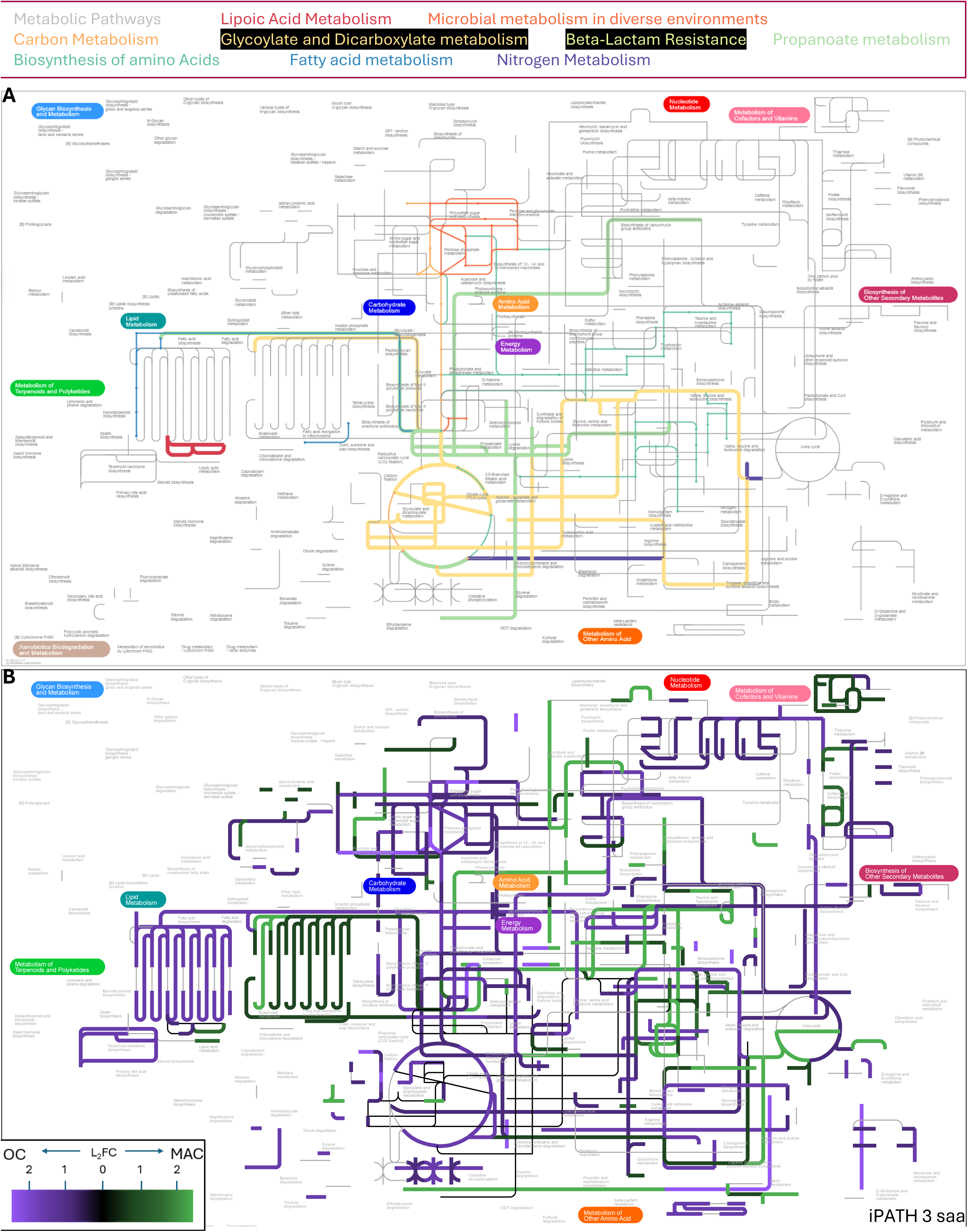
Differential expression of *S. aureus* genes in KEGG Metabolic Pathway and overlapping KEGG subterms within OCs and BMMs. **A)** An iPATH3 subway map for metabolic pathways and enriched KEGG pathways identified in Figure 1C. **B)** Differential gene expression of pathway genes for bacteria inside OCs (purple) and BMMs (green). Reactions were manually added where curation was known to be missing entirely (*aspA* and *gltA*) or partially missing (*acnA*). Maps were filtered for annotations to the *S. aureus* USA300_FPR3757 genome assembly (“saa”).

The nitrogen metabolism pathway (saa00910), which is notable for the role many of those genes play in nitrogen stress, was almost entirely upregulated by *S. aureus* inside BMMs but not OCs. For instance, TI3 inside BMMs express higher levels of *narH, narI, nirB, and nirD,* which are involved in denitrification and *agr*-dependent stimulation of the host autophagy pathway (Figure 1E) (43, 44). TI3 in BMMs also upregulate *nrdDG*, other nitrosative and oxidative stress genes that have been tested experimentally but are not annotated by KEGG (Table S1) (45–47). Interestingly, the only gene in the nitrogen metabolism pathway upregulated by *S. aureus* in OCs is *gudB* (*SAUSA300_0861*), which can also shunt nutrients into the TCA cycle (42). These data suggest *S. aureus* in OCs express less nitrosative stress than bacteria in BMMs and focus nitrogen metabolism towards generating TCA cycle metabolites.

### Osteoclasts increase metabolic gene expression and have a slower inflammatory transcriptional response to infection

We next sought to determine whether the host cell response to infection is established by cell type or time after infection. PCA of host RNA clustered samples by cell type and collection timepoint, with cell type again being the largest source of variation (Figure 3A). Analyzing host RNA-seq by comparing host cell type alone, using a Wald test as done with bacterial samples, yielded Gene Ontology (GO) terms that did not change with infection status, highlighting transcriptional changes in response to RANKL (Figure S3). Gene set enrichment analysis for Wald test DEGs showed enrichment for glycolysis and oxidative phosphorylation genes, which are known to occur during OC differentiation (Figure 3B,C) mirroring pathway expression patterns of the bacteria inside of them (Figure 2B).

**Figure 3:**
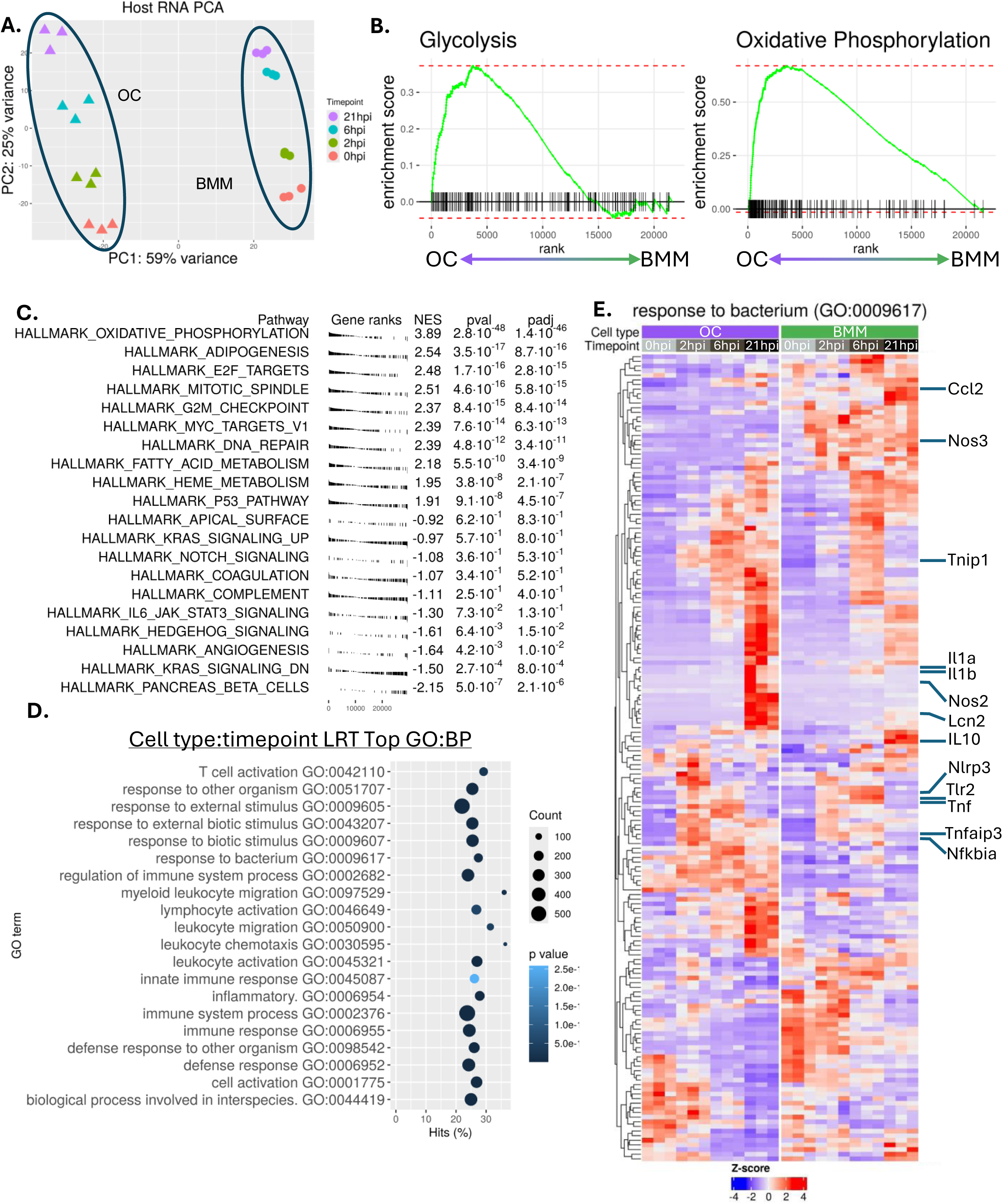
OCs express greater metabolism gene expression and respond to infection slower than BMMs. **A)** Principal component analysis of murine RNA samples 0-21hpi. **B)** Enrichment analysis of glycolytic and oxidative phosphorylation genes in OCs compared to BMMs using samples at 0hpi. **C)** Gene Set Enrichment Analysis showing the top 10 most and least enriched Hallmark pathways in OCs compared to BMMs. Comparisons for **B-C** used samples from 0-21hpi but only examined differential expression by cell type. **D)** The top 20 Gene Ontology terms enriched for genes differentially expressed by cell type over the timecourse. **E)** Heatmap for differentially expressed genes in the “Response to bacterium” GO term (GO:0009617). n=3 per timepoint, with each n representing a pool of 3 biological replicates from separate infections.

To capture the differential host cell response to infection, we switched to a likelihood-ratio test (LRT) to investigate DEGs based on the interaction of cell type and time factors. The top GO terms from LRT analysis relate to inflammation, host cell activation, and response to bacteria (Figure 3D). Examining DEG expression patterns inside the response to bacterium GO term (GO:0009617), we broadly see that by 6hpi BMMs upregulate a suite of pro-inflammatory genes, including those involved with nitric oxide synthesis (*Nos2* and *Nos3*), cytokine production (*Il1a, Il1b, Tnf, Ccl2*), and bacterial response genes (*Lcn2, Nlrp3, Tlr2*) (Figure 3E; S4E) (48–55). During the first 6hpi, OCs display greater expression of anti-inflammatory response genes (*IL10*, *Nfkbia, Tnip1, Tnfaip3*)(50, 56–61). At 21hpi, OC expression of *Il1a, Il1b,* and *Nos2* dwarfs BMM expression at any timepoint. The high expression of inflammatory genes by OCs at 21hpi, may reflect stresses from increased bacterial burden or that OCs have begun to die.

Because both OCs and *S. aureus* inside them are enriched along the TCA cycle and in genes involved with oxidative phosphorylation, we asked whether infection alters OC mitochondrial functionality. In mitochondrial stress tests we observed that infection had no impact on OC oxygen consumption rate (OCR) at 6hpi (Figure 4A). When analysis was extended to 18hpi we continued to observe no difference in OCR between mock and infected OCs (Figure S5). These data suggest infection does not impact OC mitochondrial activity or capacity.

**Figure 4:**
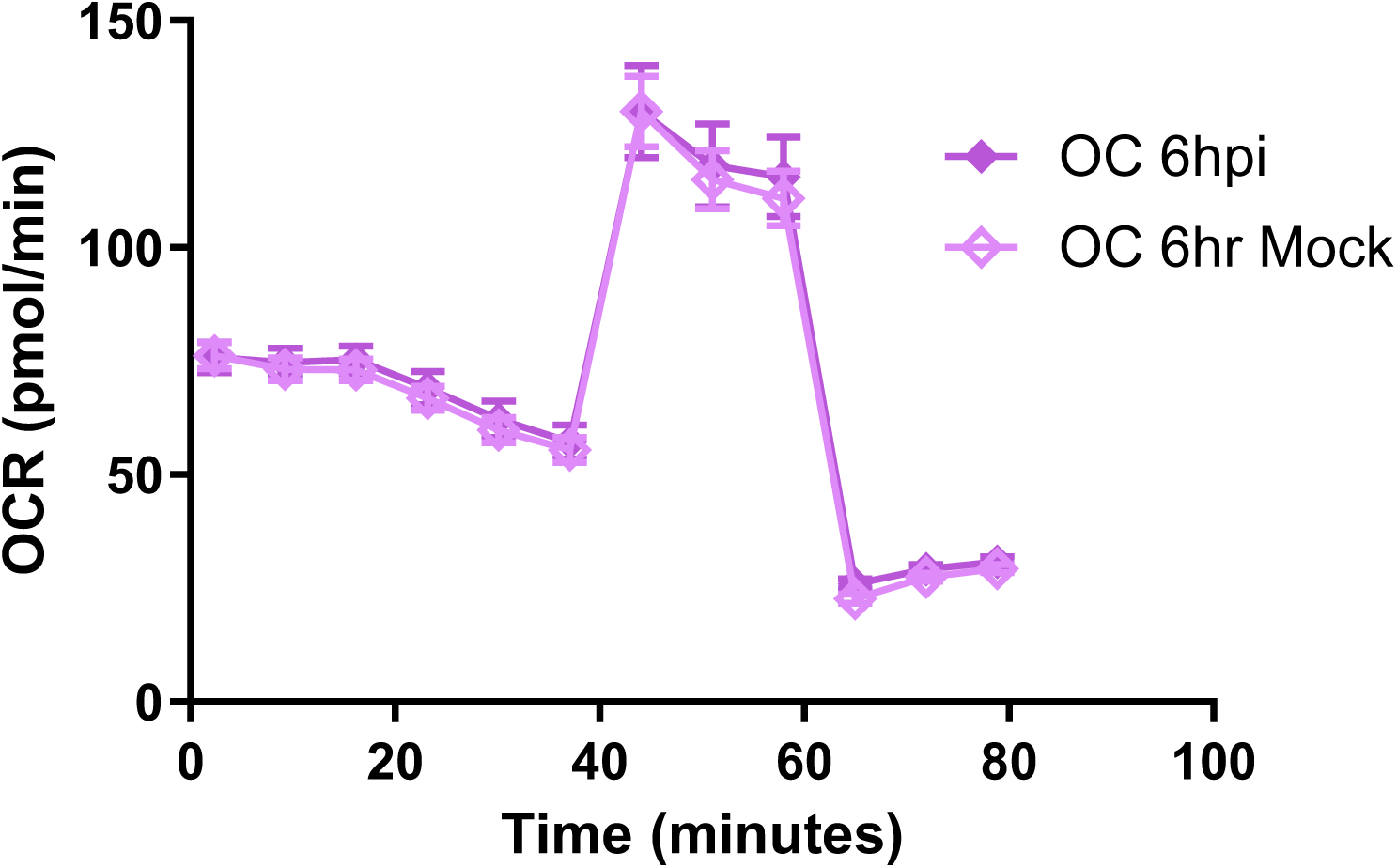
No change to mitochondrial function in OCs at 6hpi. Oxygen consumption rate data from mitochondrial stress test. Infected and mock-infected OCs show no difference at any point during the assay. Data is representative of 3 independent infections. Points are average of n= 11-12 technical replicates. Error bars indicate SEM for the experiment. p > 0.05 at all timepoints (two-way ANOVA for infection status and assay timepoint, Šídák’s correction for multiple comparisons).

### *S. aureus* relies on bacterial glycolytic genes for growth in osteoclasts

To test their role in intracellular growth, we next interrupted selected metabolic pathways by using *S. aureus* mutants. Our approach focused on acetyl-CoA, α-ketoglutarate (α-KG), and oxaloacetate (OAA) as key metabolites that serve as nodes between the TCA cycle and other metabolic processes (Figure 5A). We also focused on possible *S. aureus* nutrient sources known to be important to OCs: glucose and glutamine- which display increased uptake and flux during OC differentiation- and glutamate, which is a primary glutamine metabolite in OCs (22, 27, 28).

**Figure 5:**
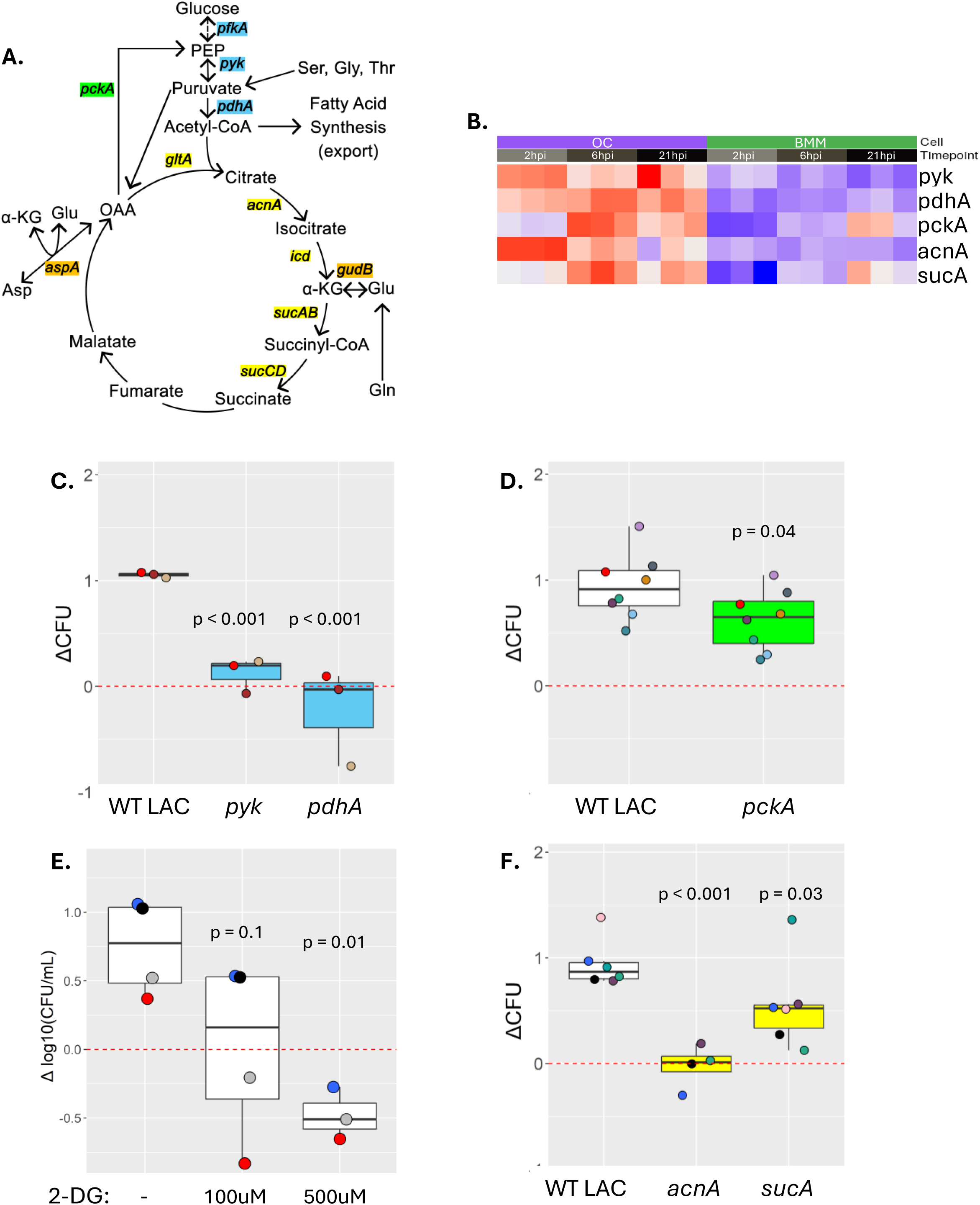
*S. aureus* requires glycolysis and *acnA* to replicate inside OCs. **A)** *S. aureus* carbon metabolism and feeder pathways. Metabolic pathways are broken down into Glycolysis and Acetyl-CoA synthesis (blue), gluconeogenesis (green), TCA cycle (yellow), and anapleurotic reactions (orange). **B)** Bacterial RNAseq heatmap for genes assessed with mutant intracellular proliferation assays. **C)** Intracellular proliferation assay of glycolysis (*pyk*), Acetyl-CoA synthesis (*pdhA*) and **D)** gluconeogenic (*pckA)* mutants examining bacterial growth between 1.5hpi and 18hpi compared to WT LAC. **E)** Addition of 2-deoxyglucose (2-DG) to OC media restricts intracellular WT LAC ΔCFU between 1.5hpi and 18hpi in a dose dependent manner. **F)** Intracellular proliferation assays of TCA cyle mutants. The *acnA* mutant had a severe growth defect and bacterial load did not change between 1.5hpi and 18hpi. The *sucA* mutant had a mild but statistically significant growth defect. Point colors represent biological replicate using same OC preparation. Statistics for **C,D,F** derived from full dataset using One-way ANOVA with Dunnett’s Test, shown in Figure S6; **E**, One-way ANOVA and Dunnett’s test.

To do so we moved from TI3 to the LAC* (AH1263) background, as it is also USA300 lineage, replicates inside OCs similarly to TI3 and has mutants readily available (8, 9). We found that across all our experiments, the bacterial load (ΔCFU) of WT LAC in OCs increases Log_10_(1.03) on average (Figure S6A). This correlates to an average 3.33 doublings of the bacteria present at 1.5hpi by the time of the 18hpi collection endpoint. All strains were inoculated with growth phase bacteria at a multiplicity of infection (MOI) 10 and internalized at similar levels (Figure S6B). As growth-phase *S. aureus* may have capacity to divide using nutrient stores obtained from liquid culture, we set an arbitrary cutoff for classification as a gene required for growth in OCs to be ΔCFU < 0.37, equating to fewer than 1.25 bacterial replications on average. Of note, we only saw one condition with reduced bacterial load at 18hpi (ΔCFU < 0), suggesting that the mutants tested generally had defects in growth but not survival when inside OCs.

We used *Δpyk::erm* to interrupt glycolysis, and *pdhA::tn* to disrupt conversion of pyruvate into acetyl-CoA. Both of these genes were upregulated by *S. aureus* inside OCs (Figure 5B). We found that OC infections with *pyk* and *pdhA* mutants resulted in severe growth impairment. The *pyk* mutant had an average ΔCFU = 0.12 (Figure 5C). The *pdhA* mutant had an average ΔCFU = -0.23. It should be noted that both *pyk* and *pdhA* grow substantially slower than WT in TSB, so inhibited mutant growth is not unique to the OC intracellular environment (Figure S8A). We attempted to extend the infection duration but were limited to <30hrs, which is when mOCs on plastic begin to spontaneously lyse, regardless of infection. We next tested necessity of gluconeogenesis using *pckA::tn*. The *pckA* mutant showed a statistically significant growth defect but was not required for intracellular growth (Figure 5D). These data suggest that glycolysis, but not gluconeogenesis, is necessary for *S. aureus* replication inside OCs.

We further investigated whether *S. aureus* requires glycolysis by using 2-deoxyglucose (2-DG) to chemically inhibit host and bacterial glycolysis. We found that WT LAC growth can be significantly blunted by 100uM 2-DG, a concentration that others found to severely reduce ATP generation with limited impairment of OC maturation (25, 62). Whereas 500uM 2-DG decreased bacterial load between 1.5 and 18hpi (Figure 5E).

### *S. aureus* utilizes TCA cycle genes to grow inside osteoclasts

We used *acnA::tn* to disrupt the TCA cycle between citrate and α-KG, and *sucA::tn* to disrupt the TCA cycle between α-KG and OAA. We found that *acnA*, which encodes the aconitase that interconverts citrate and isocitrate, is required to grow inside OCs (ΔCFU = -0.021) (Figure 5F). However, *acnA* has functions beyond TCA metabolism, and the mutant has been observed to grow slower in liquid culture (Figure S8)(63–65). Therefore, it is unclear whether the *acnA* mutant growth defect was specific to *S. aureus* in OCs or was driven by TCA cycle disruption. The *sucA* mutant had significantly blunted growth but was not required (Figure 5F). These results suggest the TCA cycle contributes to bacterial growth, but not all parts are required despite their increased expression when inside OCs.

### *S. aureus* requires aspartate transaminase but not glutamate dehydrogenase to grow in osteoclasts

We next asked whether *S. aureus* glutamine or aspartate feeder pathways to the TCA cycle were required for growth inside OCs (Figure 6A). We disrupted α-KG generation from glutamate using *gudB::tn* (42). Knocking out *gudB* resulted in a mild growth defect, with ΔCFU = 0.60 (Figure 6B). We used *aspA::tn* to disrupt transamination of Asp and α-KG into Glu and OAA(41, 42). This *aspA* mutant had a severe growth defect, with essentially no expansion inside OCs (ΔCFU = 0.140) (Figure 6C). These data suggest *S. aureus* can dispense of or compensate for *gudB* glutamate dehydrogenase activity but not the transamination activity of *aspA.* However, *aspA* functions bidirectionally, so these results leave an open question regarding whether Asp or OAA is the predominant transamination product.

**Figure 6:**
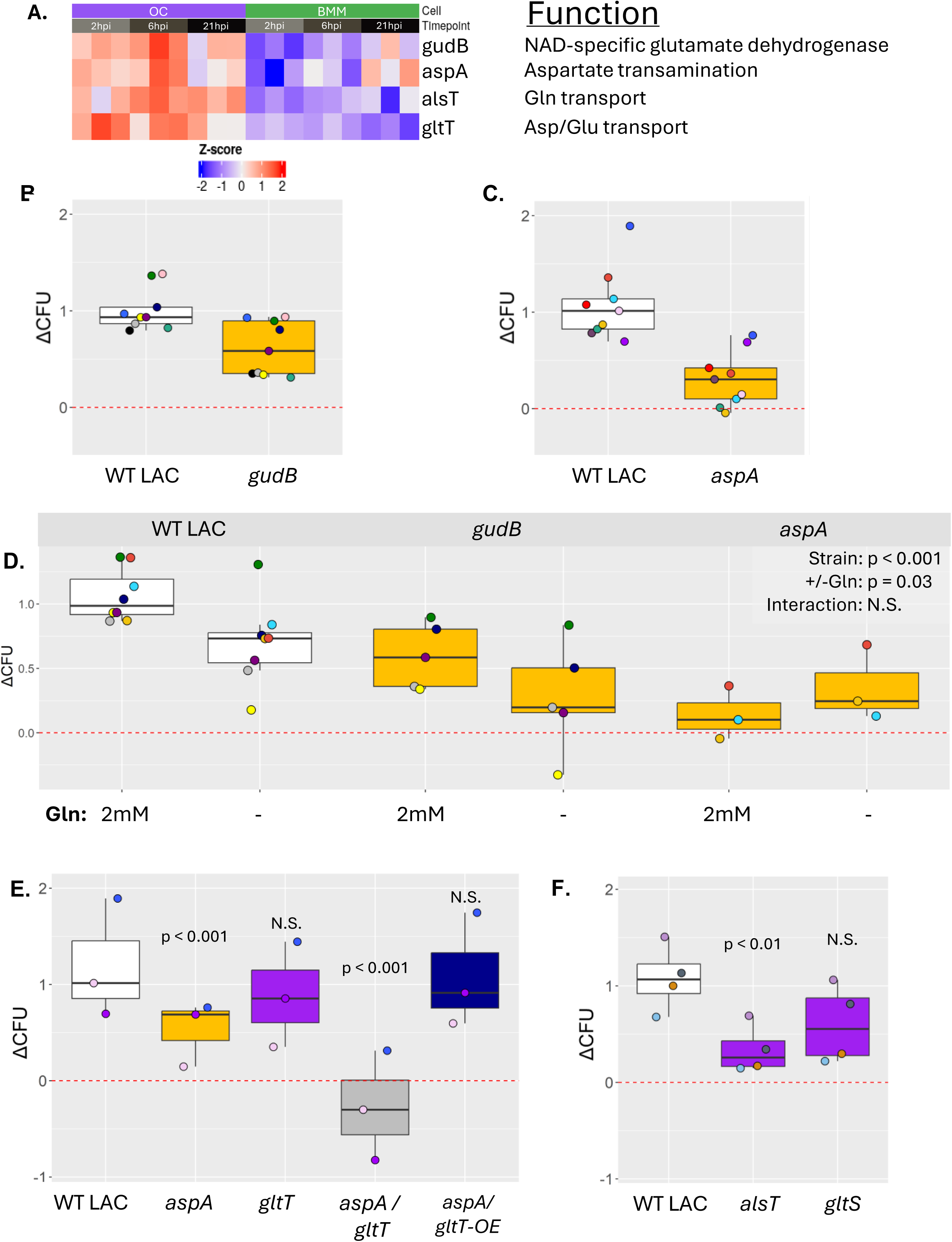
*S. aureus* utilizes Gln acquired from OCs to synthesize Asp. **A)** Bacterial RNAseq expression data for differentially expressed genes knocked out during mutant intracellular proliferation assays. The Gln transporter, *gltS*, was not differentially expressed but tested for its necessity for intracellular growth. Subsets of intracellular growth assays of mutants for **B)** glutamate dehydrogenase *gudB*, **C)** and aspartate transaminase *aspA,* **F)** Gln transporter *alsT* and Glu transporter *gltS* in comparison to WT LAC. Statistics derived from full dataset using One-way ANOVA with Dunnett’s Test, shown in Figure S6**. D)** The impact of removing Gln from the host media on intracellular growth of WT LAC, *gudB*, and *aspA.* Two-way ANOVA. **E)** Data subset of intracellular growth assays investigating the importance of Asp biosynthesis and transport using *aspA,* Asp transporter *gltT*, *aspA/gltT* double mutant, and an *aspA* mutant overexpressing gltT, with comparisons to WT LAC. Statistics derived from full dataset using One-way ANOVA with Dunnett’s Test, shown in Figure S6. Point colors represent biological replicates using same OC preparation.

We next asked whether increased metabolism of Gln to Glu and α-KG in OCs during differentiation could compensate for loss of *gudB* in *S. aureus*. We addressed this question by removing Gln from the host media, finding that WT LAC significantly reduced ΔCFU from 1.0 to 0.69, or the equivalent of 0.9 fewer bacterial doublings, while *gudB* mutant growth declined similarly, from 0.60 to 0.27 ΔCFU, a reduction of 1.1 doublings (Figure 6D). Of note, there was no synergistic effect between *gudB* knockout and Gln restriction. Since *aspA* also uses Glu and α-KG, we asked whether removing Gln from host media would impact its growth or survival, but found no change.

### *S. aureus* require *aspA-*mediated Asp biosynthesis to grow in osteoclasts

We have shown that the Asp transporter *gltT*, is inhibited by environments with a high Glu:Asp ratio (41). In such environments Asp biosynthesis via *aspA* becomes required. In mature OCs the Glu:Asp ratio is approximately 7:1, and above an inhibitory threshold for *gltT* (Figure S11)(22, 41). We therefore asked whether *aspA* is required to synthesize Asp for *S. aureus* growth in OCs, including *gltT::tn* and *aspA::tn/gltT::tn* knockouts to determine the relative importance of aspartate transport and biosynthesis. We did not observe a growth defect in the *gltT* mutant, suggesting *S. aureus* in OCs do not require Asp transport (Figure 6E). The *aspA/gltT* double mutant was the only phenotype to decrease bacterial load between 1.5hpi and 18hpi in OCs with ΔCFU = - 0.27. We then used a strain overexpressing *gltT* on the *aspA* knockout background to overcome glutamate inhibition of Asp transport, finding a complete rescue of bacterial growth inside OCs (ΔCFU = 1.09). These data, coupled with *aspA* mutant infection data, show that Asp metabolism is vital for *S. aureus* growth in OCs.

### *S. aureus* acquire Gln from osteoclasts to grow intracellularly

Our data showed that restricting Gln in the OC media blunts *S. aureus* intracellular growth. While OCs import greater amounts of Gln than other metabolites, it is quickly metabolized into Glu, α-KG, and several other metabolites not observed in their precursors (22). Restricting Gln to OCs, therefore, lowers availability of numerous metabolites. We sought to identify whether Gln or Glu are the metabolites imported by *S. aureus* using mutants *alsT::tn* and *gltS::tn*, which disrupt the primary transporters of Gln and Glu, respectively (66). The *alsT* mutant had a severe growth defect, with ΔCFU = 0.34, suggesting Gln transport is required for growth in OCs (Figure 6F). Meanwhile growth of the *gltS* mutant did not significantly differ from WT. These experiments show that intracellular *S. aureus* require Gln but not Glu transport to grow inside OCs.

## DISCUSSION

*S. aureus* uses many tactics to avoid killing by the immune system and antibiotics, including intracellular infection. We have previously shown that *S. aureus* can rapidly expand in OCs, suggesting that OCs may be a reservoir for persistent or recurrent infection. In this study we sought to identify how *S. aureus* are able to expand in OCs, but not their BMM precursors. Through infections using *S. aureus* metabolism and transport mutants and manipulating OC culture conditions we identified that *S. aureus* growth in OCs is dependent on OC access to Gln and its own ability to transport it. *S. aureus* growth in OCs is also dependent on their own glycolytic capability. While we found several conditions that abrogated bacterial growth in OCs, we found only one that resulted in a meaningful decrease in bacterial load overnight. Therefore, the increased intracellular bacterial load appears to result from active provision of necessary nutrients as well as lack of an effective antimicrobial response in OCs.

Using RNA sequencing, we observed that *S. aureus* in OCs increase expression of metabolism genes compared to bacteria in BMMs. This was unsurprising, as *S. aureus* has been shown to access nutrients phagocytosed by their host and OCs increase glucose and Gln uptake during differentiation (22, 67, 68). Although Gln is metabolized into many compounds by OCs, the severe growth defect observed in the *alsT* mutant suggests that Gln, rather than OC-produced Gln metabolites, is a primary nutrient fueling intracellular growth.

Many metabolic genes were upregulated in bacteria in OCs. We tested *pyk*, *pdhA*, *acnA*, and *aspA* and found that they contribute significantly to bacterial growth in OCs at the analyzed timepoints. Previous studies have shown *pyk, pdhA,* and *acnA* are important for growth even in liquid culture conditions (63, 69, 70). Therefore, growth defects observed inside OCs may reflect baseline changes to *S. aureus* growth kinetics when glycolysis, acetyl-CoA synthesis, or aconitase activity is disrupted. In contrast, the *aspA* mutant grew in liquid culture at similar rates as *gudB, pckA,* and *aspA/*pOS1*-gltT*, suggesting the *aspA* mutant growth defect inside OCs is not due to an underlying growth defect.

We previously found that *S. aureus* in media with a high Glu: Asp ratio rely on *aspA* for Asp biosynthesis to compensate for inhibition of *gltT* mediated Asp transport (41). Data from Hu *et al* show that OCs and their precursors have Glu:Asp ratios of at least 4:1 (Figure S11) (22). Those data suggest *gltT* may be inhibited in BMMs and OCs. In the previous work and here, *gltT* overexpression compensated for *aspA* mutants, suggesting that *S. aureus* growth is limited by Asp availability both in bone and within OCs. The importance of Asp availability appears to be emphasized by the fact that the *aspA/gltT* double mutant was the only strain to have a significant reduction in viable bacteria inside OCs at 18hpi. We suspect this reduced viability is because the mutant cannot acquire enough Asp to survive. However, we cannot currently exclude the possibility of active OC clearance of the bacteria.

All but one of the *S. aureus* mutant strains tested had ΔCFU greater than 0. This suggests that mutant bacteria in OCs were able to survive, even though they did not proliferate. Our RNA-seq data show that OCs have blunted or delayed transcriptional response to infection, while *S. aureus* in OCs express fewer genes associated with nitrosative stress. In BMMs, mounting an oxidative burst is important for bacterial killing and can induce several of the *S. aureus* nitrosative stress genes that were differentially expressed between BMMs and OCs (46, 71, 72). These data suggest that OCs do not have an effective defense response and may not mount an oxidative burst. Supporting this, disrupting glycolysis, which is required to survive nitrosative stress(45), did not lead to decreased survival of a pyk mutant in OCs.

In various macrophage populations, the response to infection can be altered by exposure to inflammatory factors, like IL-4 or RANKL (8, 73–75). While RANKL may blunt the cell’s defense response independent of OC differentiation, it is unclear how cellular changes related to OC maturation may also impact the cell’s ability to fight *S. aureus* infection. As such, further investigation of the OC response to infection, or lack thereof, is warranted.

While our experiments produced robust results, our study has several limitations. First, our RNA-seq dataset does not have a timecourse of mock infection. OCs continue to mature during post-infection incubation, and at an accelerated rate compared to uninfected cells. Therefore, it is difficult to directly match degree of differentiation in primary cell cultures. Second, the primary cells used in our study can differentiate at variable rates. In our experiments with mutants and media manipulations, we controlled for this by including a WT LAC condition for each OC cell preparation as a technical control. However, biological replicates with different OC cultures include OCs at slightly different stages of maturity. Third, our study examines infection dynamics of OCs plated on plastic and others have shown that the metabolic profile differs for OCs that are actively resorbing (27). While metabolic differences between plastic and native bone may change some metabolic requirements for intracellular *S. aureus*, Asp biosynthesis has already been found to be important for bacterial survival within bone in vivo (41).

In conclusion, OCs are permissive hosts for S. aureus due to nutrient abundance and deficiencies in their bacterial defense response. Aspartate biosynthesis and import of host Gln each contribute to growth of these bacteria in OCs. These metabolic host-pathogen interactions may provide novel opportunities to address intracellular reservoirs contributing to osteomyelitis pathogenesis.

## ACKNOWLEDGEMENTS

We would like to thank the Integrative Informatics Core of the WashU Rheumatic Diseases Research Resource-based Center (NIH P30-AR073752) for assistance in experimental design and analyses for this manuscript. We thank the Diabetes research Center (NIH P30-DK020579) for the use of the Seahorse instrument. We thank the Genome Technology Access Center at the McDonnell Genome Institute at Washington University School of Medicine for help with genomic analysis. GTAC is partially supported P30-CA91842. We thank the Richardson Lab, who provided the *Δpyk::erm* LAC mutant. We would like to thank Linda Cox for her work maintaining mouse colonies used in this study.

## FUNDING

LDO was supported by the NIH grant R01AI161022, the Training Program in Cellular and Molecular Biology, T32GM139774, and Skeletal Disorders Training Program, T32AR060719. SB was supported by NIH grant R01AI161022. NR was supported by NIH grants R01CA258325 and P30AR073752. GM was supported by NIH grants R01AI161022, and R01AG077732. JEC was supported by NIH grants R01AI173795, R01AI177615, and R01AI161022. DJV was supported by NIH grants R01AI161022, R21EB034882, and P30AR074992.

## References

1. Hofstee MI, Muthukrishnan G, Atkins GJ, Riool M, Thompson K, Morgenstern M, Stoddart MJ, Richards RG, Zaat SAJ, Moriarty TF. 2020. Current Concepts of Osteomyelitis: From Pathologic Mechanisms to Advanced Research Methods. Am J Pathol 190:1151–1163.

2. Conterno LO, Turchi MD. 2013. Antibiotics for treating chronic osteomyelitis in adults. Cochrane Database Syst Rev CD004439.

3. Gimza BD, Cassat JE. 2021. Mechanisms of Antibiotic Failure During Staphylococcus aureus Osteomyelitis. Front Immunol 12.

4. Yang D, Wijenayaka AR, Solomon LB, Pederson SM, Findlay DM, Kidd SP, Atkins GJ. 2018. Novel Insights into Staphylococcus aureus Deep Bone Infections: the Involvement of Osteocytes. mBio 9:e00415–18.

5. Granata V, Possetti V, Parente R, Bottazzi B, Inforzato A, Sobacchi C. 2022. The osteoblast secretome in Staphylococcus aureus osteomyelitis. Frontiers in Immunology 13.

6. Josse J, Velard F, Gangloff SC. 2015. Staphylococcus aureus vs. Osteoblast: Relationship and Consequences in Osteomyelitis. Frontiers in Cellular and Infection Microbiology 5.

7. Trouillet-Assant S, Gallet M, Nauroy P, Rasigade J-P, Flammier S, Parroche P, Marvel J, Ferry T, Vandenesch F, Jurdic P, Laurent F. 2015. Dual Impact of Live Staphylococcus aureus on the Osteoclast Lineage, Leading to Increased Bone Resorption. The Journal of Infectious Diseases 211:571–581.

8. Krauss JL, Roper PM, Ballard A, Shih C-C, Fitzpatrick JAJ, Cassat JE, Ng PY, Pavlos NJ, Veis DJ. 2019. Staphylococcus aureus Infects Osteoclasts and Replicates Intracellularly. mBio 10:e02447–19.

9. Roper PM, Eichelberger KR, Cox L, O’Connor L, Shao C, Ford CA, Fritz SA, Cassat JE, Veis DJ. 2021. Contemporary Clinical Isolates of Staphylococcus aureus from Pediatric Osteomyelitis Patients Display Unique Characteristics in a Mouse Model of Hematogenous Osteomyelitis. Infect Immun 89:e0018021.

10. Veis DJ, O’Brien CA. 2023. Osteoclasts, Master Sculptors of Bone. Annu Rev Pathol Mech Dis 18:257–281.

11. McDonald MM, Kim AS, Mulholland BS, Rauner M. 2021. New Insights Into Osteoclast Biology. JBMR Plus 5:e10539.

12. Marks SC, Seifert MF. 1985. The lifespan of osteoclasts: Experimental studies using the giant granule cytoplasmic marker characteristic of beige mice. Bone 6:451–455.

13. Loutit JF, Townsend KM. 1982. Longevity of osteoclasts in radiation chimaeras of osteopetrotic beige and normal mice. Br J Exp Pathol 63:221–223.

14. Marshall MJ, Davie MWJ. 1991. An immunocytochemical method for studying the kinetics of osteoclast nuclei on intact mouse parietal bone. Histochem J 23:402–408.

15. Young RW. 1962. CELL PROLIFERATION AND SPECIALIZATION DURING ENDOCHONDRAL OSTEOGENESIS IN YOUNG RATS. The Journal of Cell Biology 14:357–370.

16. Jacome-Galarza CE, Percin GI, Muller JT, Mass E, Lazarov T, Eitler J, Rauner M, Yadav VK, Crozet L, Bohm M, Loyher P-L, Karsenty G, Waskow C, Geissmann F. 2019. Developmental origin, functional maintenance and genetic rescue of osteoclasts. Nature 568:541–545.

17. Yahara Y, Barrientos T, Tang YJ, Puviindran V, Nadesan P, Zhang H, Gibson JR, Gregory SG, Diao Y, Xiang Y, Qadri YJ, Souma T, Shinohara ML, Alman BA. 2020. Erythromyeloid progenitors give rise to a population of osteoclasts that contribute to bone homeostasis and repair. Nat Cell Biol 22:49–59.

18. Petronglo JR, Putnam NE, Ford CA, Cruz-Victorio V, Curry JM, Butrico CE, Fulbright LE, Johnson JR, Peck SH, Fatah SR, Cassat JE. 2022. Context-Dependent Roles for Toll-Like Receptors 2 and 9 in the Pathogenesis of Staphylococcus aureus Osteomyelitis. Infection and Immunity 0:e00417–22.

19. Chen Z, Su L, Xu Q, Katz J, Michalek SM, Fan M, Feng X, Zhang P. 2015. IL-1R/TLR2 through MyD88 Divergently Modulates Osteoclastogenesis through Regulation of Nuclear Factor of Activated T Cells c1 (NFATc1) and B Lymphocyte-induced Maturation Protein-1 (Blimp1). J Biol Chem 290:30163–30174.

20. Sun Y, Li J, Xie X, Gu F, Sui Z, Zhang K, Yu T. 2021. Macrophage-Osteoclast Associations: Origin, Polarization, and Subgroups. Front Immunol 12:778078.

21. Das BK, Wang L, Fujiwara T, Zhou J, Aykin-Burns N, Krager KJ, Lan R, Mackintosh SG, Edmondson R, Jennings ML, Wang X, Feng JQ, Barrientos T, Gogoi J, Kannan A, Gao L, Xing W, Mohan S, Zhao H. 2022. Transferrin receptor 1-mediated iron uptake regulates bone mass in mice via osteoclast mitochondria and cytoskeleton. eLife 11:e73539.

22. Hu G, Yu Y, Ren Y, Tower RJ, Zhang G-F, Karner CM. 2024. Glutaminolysis provides nucleotides and amino acids to regulate osteoclast differentiation in mice. EMBO Rep 25:4515–4541.

23. Ishii K, Fumoto T, Iwai K, Takeshita S, Ito M, Shimohata N, Aburatani H, Taketani S, Lelliott CJ, Vidal-Puig A, Ikeda K. 2009. Coordination of PGC-1β and iron uptake in mitochondrial biogenesis and osteoclast activation. Nat Med 15:259–266.

24. Zeng R, Faccio R, Novack DV. 2015. Alternative NF-κB Regulates RANKL-Induced Osteoclast Differentiation and Mitochondrial Biogenesis via Independent Mechanisms. Journal of Bone and Mineral Research 30:2287–2299.

25. Li B, Lee W-C, Song C, Ye L, Abel ED, Long F. 2020. Both aerobic glycolysis and mitochondrial respiration are required for osteoclast differentiation. The FASEB Journal 34:11058–11067.

26. Peng R, Dong Y, Zheng M, Kang H, Wang P, Zhu M, Song K, Wu W, Li F. 2024. IL-17 promotes osteoclast-induced bone loss by regulating glutamine-dependent energy metabolism. Cell Death Dis 15:111.

27. Lemma S, Sboarina M, Porporato PE, Zini N, Sonveaux P, Di Pompo G, Baldini N, Avnet S. 2016. Energy metabolism in osteoclast formation and activity. The International Journal of Biochemistry & Cell Biology 79:168–180.

28. Indo Y, Takeshita S, Ishii K-A, Hoshii T, Aburatani H, Hirao A, Ikeda K. 2013. Metabolic regulation of osteoclast differentiation and function. Journal of Bone and Mineral Research 28:2392–2399.

29. Takeshita S, Kaji K, Kudo A. 2000. Identification and Characterization of the New Osteoclast Progenitor with Macrophage Phenotypes Being Able to Differentiate into Mature Osteoclasts*. J Bone Miner Res 15:1477–1488.

30. McHugh KP, Hodivala-Dilke K, Zheng M-H, Namba N, Lam J, Novack D, Feng X, Ross FP, Hynes RO, Teitelbaum SL. 2000. Mice lacking β3 integrins are osteosclerotic because of dysfunctional osteoclasts. J Clin Invest 105:433–440.

31. Pisu D, Huang L, Rin Lee BN, Grenier JK, Russell DG. 2020. Dual RNA-Sequencing of *Mycobacterium tuberculosis*-Infected Cells from a Murine Infection Model. STAR Protocols 1:100123.

32. Kim D, Paggi JM, Park C, Bennett C, Salzberg SL. 2019. Graph-based genome alignment and genotyping with HISAT2 and HISAT-genotype. Nat Biotechnol 37:907–915.

33. Liao Y, Smyth GK, Shi W. 2019. The R package Rsubread is easier, faster, cheaper and better for alignment and quantification of RNA sequencing reads. Nucleic Acids Res 47:e47.

34. Liao Y, Smyth GK, Shi W. 2014. featureCounts: an efficient general purpose program for assigning sequence reads to genomic features. Bioinformatics 30:923–930.

35. Chisanga D, Liao Y, Shi W. 2022. Impact of gene annotation choice on the quantification of RNA-seq data. BMC Bioinformatics 23:107.

36. Love MI, Huber W, Anders S. 2014. Moderated estimation of fold change and dispersion for RNA-seq data with DESeq2. Genome Biology 15:550.

37. Martin FJ, Amode MR, Aneja A, Austine-Orimoloye O, Azov AG, Barnes I, Becker A, Bennett R, Berry A, Bhai J, Bhurji SK, Bignell A, Boddu S, Branco Lins PR, Brooks L, Ramaraju SB, Charkhchi M, Cockburn A, Da Rin Fiorretto L, Davidson C, Dodiya K, Donaldson S, El Houdaigui B, El Naboulsi T, Fatima R, Giron CG, Genez T, Ghattaoraya GS, Martinez JG, Guijarro C, Hardy M, Hollis Z, Hourlier T, Hunt T, Kay M, Kaykala V, Le T, Lemos D, Marques-Coelho D, Marugán JC, Merino GA, Mirabueno LP, Mushtaq A, Hossain SN, Ogeh DN, Sakthivel MP, Parker A, Perry M, Piližota I, Prosovetskaia I, Pérez-Silva JG, Salam AIA, Saraiva-Agostinho N, Schuilenburg H, Sheppard D, Sinha S, Sipos B, Stark W, Steed E, Sukumaran R, Sumathipala D, Suner M-M, Surapaneni L, Sutinen K, Szpak M, Tricomi FF, Urbina-Gómez D, Veidenberg A, Walsh TA, Walts B, Wass E, Willhoft N, Allen J, Alvarez-Jarreta J, Chakiachvili M, Flint B, Giorgetti S, Haggerty L, Ilsley GR, Loveland JE, Moore B, Mudge JM, Tate J, Thybert D, Trevanion SJ, Winterbottom A, Frankish A, Hunt SE, Ruffier M, Cunningham F, Dyer S, Finn RD, Howe KL, Harrison PW, Yates AD, Flicek P. 2023. Ensembl 2023. Nucleic Acids Res 51:D933–D941.

38. Diep BA, Gill SR, Chang RF, Phan TH, Chen JH, Davidson MG, Lin F, Lin J, Carleton HA, Mongodin EF, Sensabaugh GF, Perdreau-Remington F. 2006. Complete genome sequence of USA300, an epidemic clone of community-acquired meticillin-resistant *Staphylococcus aureus*. The Lancet 367:731–739.

39. Rohatgi N, Zou W, Li Y, Cho K, Collins PL, Tycksen E, Pandey G, DeSelm CJ, Patti GJ, Dey A, Teitelbaum SL. 2023. BAP1 promotes osteoclast function by metabolic reprogramming. Nat Commun 14:5923.

40. Roper PM, Beetner J, Davis JL, Shao C, Zhang J, Cox L, Pokhrel NK, Cassat JE, Rangel-Moreno J, Muthukrishnan G, Veis DJ. 2025. Characterization of a mouse model of osteoarticular infection using clinical isolates of Staphylococcus aureus. JBMR Plus 9:ziaf093.

41. Potter AD, Butrico CE, Ford CA, Curry JM, Trenary IA, Tummarakota SS, Hendrix AS, Young JD, Cassat JE. 2020. Host nutrient milieu drives an essential role for aspartate biosynthesis during invasive Staphylococcus aureus infection. Proceedings of the National Academy of Sciences 117:12394–12401.

42. Halsey CR, Lei S, Wax JK, Lehman MK, Nuxoll AS, Steinke L, Sadykov M, Powers R, Fey PD. 2017. Amino Acid Catabolism in Staphylococcus aureus and the Function of Carbon Catabolite Repression. mBio 8:e01434–16.

43. Nurxat N, Wang Q, Zhao N, Guo Y, Zhang X, Wang Y, Jian Y, Wang H, Yang S, Li M, Liu Q. 2025. Endogenous nitric oxide promotes Staphylococcus aureus virulence by activating autophagy. mBio 16:e04006–24.

44. Li Y, Pan T, Cao R, Li W, He Z, Sun B. 2023. Nitrate Reductase NarGHJI Modulates Virulence via Regulation of agr Expression in Methicillin-Resistant Staphylococcus aureus Strain USA300 LAC. Microbiology Spectrum 11:e03596–22.

45. Grosser MR, Paluscio E, Thurlow LR, Dillon MM, Cooper VS, Kawula TH, Richardson AR. 2018. Genetic requirements for Staphylococcus aureus nitric oxide resistance and virulence. PLOS Pathogens 14:e1006907.

46. Richardson AR, Dunman PM, Fang FC. 2006. The nitrosative stress response of Staphylococcus aureus is required for resistance to innate immunity. Molecular Microbiology 61:927–939.

47. Chang W, Small DA, Toghrol F, Bentley WE. 2006. Global Transcriptome Analysis of Staphylococcus aureus Response to Hydrogen Peroxide. Journal of Bacteriology 188:1648–1659.

48. Bailey JD, Diotallevi M, Nicol T, McNeill E, Shaw A, Chuaiphichai S, Hale A, Starr A, Nandi M, Stylianou E, McShane H, Davis S, Fischer R, Kessler BM, McCullagh J, Channon KM, Crabtree MJ. 2019. Nitric Oxide Modulates Metabolic Remodeling in Inflammatory Macrophages through TCA Cycle Regulation and Itaconate Accumulation. Cell Reports 28:218–230.e7.

49. Connelly L, Jacobs AT, Palacios-Callender M, Moncada S, Hobbs AJ. 2003. Macrophage Endothelial Nitric-oxide Synthase Autoregulates Cellular Activation and Pro-inflammatory Protein Expression*. Journal of Biological Chemistry 278:26480–26487.

50. Liu T, Zhang L, Joo D, Sun S-C. 2017. NF-κB signaling in inflammation. Sig Transduct Target Ther 2:17023.

51. Toyonaga T, Matsuura M, Mori K, Honzawa Y, Minami N, Yamada S, Kobayashi T, Hibi T, Nakase H. 2016. Lipocalin 2 prevents intestinal inflammation by enhancing phagocytic bacterial clearance in macrophages. Sci Rep 6:35014.

52. Wang Q, Li S, Tang X, Liang L, Wang F, Du H. 2019. Lipocalin 2 Protects Against Escherichia coli Infection by Modulating Neutrophil and Macrophage Function. Front Immunol 10:2594.

53. Flo TH, Smith KD, Sato S, Rodriguez DJ, Holmes MA, Strong RK, Akira S, Aderem A. 2004. Lipocalin 2 mediates an innate immune response to bacterial infection by sequestrating iron. Nature 432:917–921.

54. El-Zayat SR, Sibaii H, Mannaa FA. 2019. Toll-like receptors activation, signaling, and targeting: an overview. Bulletin of the National Research Centre 43:187.

55. Gschwandtner M, Derler R, Midwood KS. 2019. More Than Just Attractive: How CCL2 Influences Myeloid Cell Behavior Beyond Chemotaxis. Front Immunol 10.

56. Iyer SS, Cheng G. 2012. Role of Interleukin 10 Transcriptional Regulation in Inflammation and Autoimmune Disease. Crit Rev Immunol 32:23–63.

57. Mauro C, Pacifico F, Lavorgna A, Mellone S, Iannetti A, Acquaviva R, Formisano S, Vito P, Leonardi A. 2006. ABIN-1 Binds to NEMO/IKKγ and Co-operates with A20 in Inhibiting NF-κB*. Journal of Biological Chemistry 281:18482–18488.

58. G’Sell RT, Gaffney PM, Powell DW. 2015. Review: A20-Binding Inhibitor of NF-κB Activation 1 Is a Physiologic Inhibitor of NF-κB: A Molecular Switch for Inflammation and Autoimmunity. Arthritis & Rheumatology 67:2292–2302.

59. Das T, Chen Z, Hendriks RW, Kool M. 2018. A20/Tumor Necrosis Factor α-Induced Protein 3 in Immune Cells Controls Development of Autoinflammation and Autoimmunity: Lessons from Mouse Models. Front Immunol 9.

60. Wertz IE, O’Rourke KM, Zhou H, Eby M, Aravind L, Seshagiri S, Wu P, Wiesmann C, Baker R, Boone DL, Ma A, Koonin EV, Dixit VM. 2004. De-ubiquitination and ubiquitin ligase domains of A20 downregulate NF-κB signalling. Nature 430:694–699.

61. Baima ET, Guzova JA, Mathialagan S, Nagiec EE, Hardy MM, Song LR, Bonar SL, Weinberg RA, Selness SR, Woodard SS, Chrencik J, Hood WF, Schindler JF, Kishore N, Mbalaviele G. 2010. Novel Insights into the Cellular Mechanisms of the Anti-inflammatory Effects of NF-κB Essential Modulator Binding Domain Peptides. Journal of Biological Chemistry 285:13498–13506.

62. Murata K, Fang C, Terao C, Giannopoulou EG, Lee YJ, Lee MJ, Mun S-H, Bae S, Qiao Y, Yuan R, Furu M, Ito H, Ohmura K, Matsuda S, Mimori T, Matsuda F, Park-Min K-H, Ivashkiv LB. 2017. Hypoxia-Sensitive COMMD1 Integrates Signaling and Cellular Metabolism in Human Macrophages and Suppresses Osteoclastogenesis. Immunity 47:66–79.e5.

63. Somerville GA, Chaussee MS, Morgan CI, Fitzgerald JR, Dorward DW, Reitzer LJ, Musser JM. 2002. Staphylococcus aureus Aconitase Inactivation Unexpectedly Inhibits Post-Exponential-Phase Growth and Enhances Stationary-Phase Survival. Infect Immun 70:6373–6382.

64. Chen F, Zhao Q, Yang Z, Chen R, Pan H, Wang Y, Liu H, Cao Q, Gan J, Liu X, Zhang N, Yang C-G, Liang H, Lan L. 2024. Citrate serves as a signal molecule to modulate carbon metabolism and iron homeostasis in Staphylococcus aureus. PLoS Pathog 20:e1012425.

65. Ding Y, Liu X, Chen F, Di H, Xu B, Zhou L, Deng X, Wu M, Yang C-G, Lan L. 2014. Metabolic sensor governing bacterial virulence in Staphylococcus aureus. Proc Natl Acad Sci U S A 111:E4981–E4990.

66. Zeden MS, Kviatkovski I, Schuster CF, Thomas VC, Fey PD, Gründling A. 2020. Identification of the main glutamine and glutamate transporters in Staphylococcus aureus and their impact on c-di-AMP production. Mol Microbiol 113:1085–1100.

67. Flannagan RS, Heinrichs DE. 2020. Macrophage-driven nutrient delivery to phagosomal Staphylococcus aureus supports bacterial growth. EMBO Rep 21:EMBR202050348.

68. Arnett TR, Orriss IR. 2018. Metabolic properties of the osteoclast. Bone 115:25–30.

69. Vitko NP, Spahich NA, Richardson AR. 2015. Glycolytic Dependency of High-Level Nitric Oxide Resistance and Virulence in Staphylococcus aureus. mBio 6:10.1128/mbio.00045-15.

70. Kim G-L, Hooven TA, Norambuena J, Li B, Boyd JM, Yang JH, Parker D. 2021. Growth and Stress Tolerance Comprise Independent Metabolic Strategies Critical for Staphylococcus aureus Infection. mBio 12:10.1128/mbio.00814-21.

71. Dupré-Crochet S, Erard M, Nüβe O. 2013. ROS production in phagocytes: why, when, and where? J Leukoc Biol 94:657–670.

72. Pidwill GR, Gibson JF, Cole J, Renshaw SA, Foster SJ. 2021. The Role of Macrophages in Staphylococcus aureus Infection. Front Immunol 11:620339.

73. Siwczak F, Cseresnyes Z, Hassan MIA, Aina KO, Carlstedt S, Sigmund A, Groger M, Surewaard BGJ, Werz O, Figge MT, Tuchscherr L, Loffler B, Mosig AS. 2022. Human macrophage polarization determines bacterial persistence of Staphylococcus aureus in a liver-on-chip-based infection model. Biomaterials 287:121632.

74. Werz O, Gerstmeier J, Libreros S, De la Rosa X, Werner M, Norris PC, Chiang N, Serhan CN. 2018. Human macrophages differentially produce specific resolvin or leukotriene signals that depend on bacterial pathogenicity. Nat Commun 9:59.

75. Mota RF, Cavalcanti de Araújo PH, Cezine MER, Matsuo FS, Metzner RJM, Oliveira de Biagi Junior CA, Peronni KC, Hayashi H, Shimamura M, Nakagami H, Osako MK. 2022. RANKL Impairs the TLR4 Pathway by Increasing TRAF6 and RANK Interaction in Macrophages. Biomed Res Int 2022:7740079.

76. Boles BR, Thoendel M, Roth AJ, Horswill AR. 2010. Identification of Genes Involved in Polysaccharide-Independent Staphylococcus aureus Biofilm Formation. PLOS ONE 5:e10146.

77. Fey PD, Endres JL, Yajjala VK, Widhelm TJ, Boissy RJ, Bose JL, Bayles KW. 2013. A genetic resource for rapid and comprehensive phenotype screening of nonessential Staphylococcus aureus genes. mBio 4:e00537–00512.

